# Impaired development of the medial olivocochlear system in a KCNQ4-deficient mouse model

**DOI:** 10.64898/2026.01.21.700803

**Authors:** Ezequiel Rías, Ingrid Ouwerkerk, Guillermo Spitzmaul, Leonardo Dionisio

**Affiliations:** Instituto de Investigaciones Bioquímicas de Bahía Blanca (INIBIBB), CONICET-UNS, Bahía Blanca, Argentina; Departamento de Biología, Bioquímica y Farmacia (BByF), UNS, Bahía Blanca, Argentina

**Keywords:** Outer Hair Cell, Efferent terminals, KCNQ4, Inner ear

## Abstract

The medial olivocochlear (MOC) efferent system modulates outer hair cell (OHC) excitability and protects cochlea from overstimulation. Cholinergic activation of α9α10 nicotinic acetylcholine receptors (nAChRs) triggers Ca⁺² influx, activating BK and SK2 Ca⁺²-dependent K⁺ channels, and K⁺ extrusion through KCNQ4 to restore membrane potential. KCNQ4-loss causes chronic depolarization, OHC dysfunction, and hearing loss. Here, we investigated how KCNQ4 deficiency affects cochlear efferent synapse development and organization.

Using confocal immunofluorescence, we analyzed efferent innervation in the organ of Corti of *Kcnq4^−/−^* (KO) and *Kcnq4^+/+^*(WT) mice at 2, 3, 4, and 10 postnatal weeks (W). At 2 W, efferent terminals were similarly distributed between basal and lateral OHC membrane domains in both genotypes. During maturation, WT mice exhibited complete relocation of MOC terminals to the basal domain, whereas KO mice showed delayed maturation, with some terminals laterally displaced up to 10 W. KCNQ4 absence was associated with reduced number and volume of efferent boutons on OHCs. Milder morphometric alterations were observed in efferent boutons within the inner hair cell region.

At the molecular level, qPCR revealed downregulation of α10 nAChR subunit, BK, and SK2 transcripts in KO at 4 W, with recovery to 10 W. Despite this recovery, BK protein showed reduced expression, mislocalization, and disorganized synaptic plaques in OHCs. KO also displayed age-dependent upregulation of the calcium-binding proteins calbindin and calretinin, suggesting compensatory responses to altered Ca^+^² homeostasis.

Together, these findings demonstrate that KCNQ4 is essential for OHC repolarization, maturation and maintenance of cochlear efferent synapses.

**Graphical Abstract:** 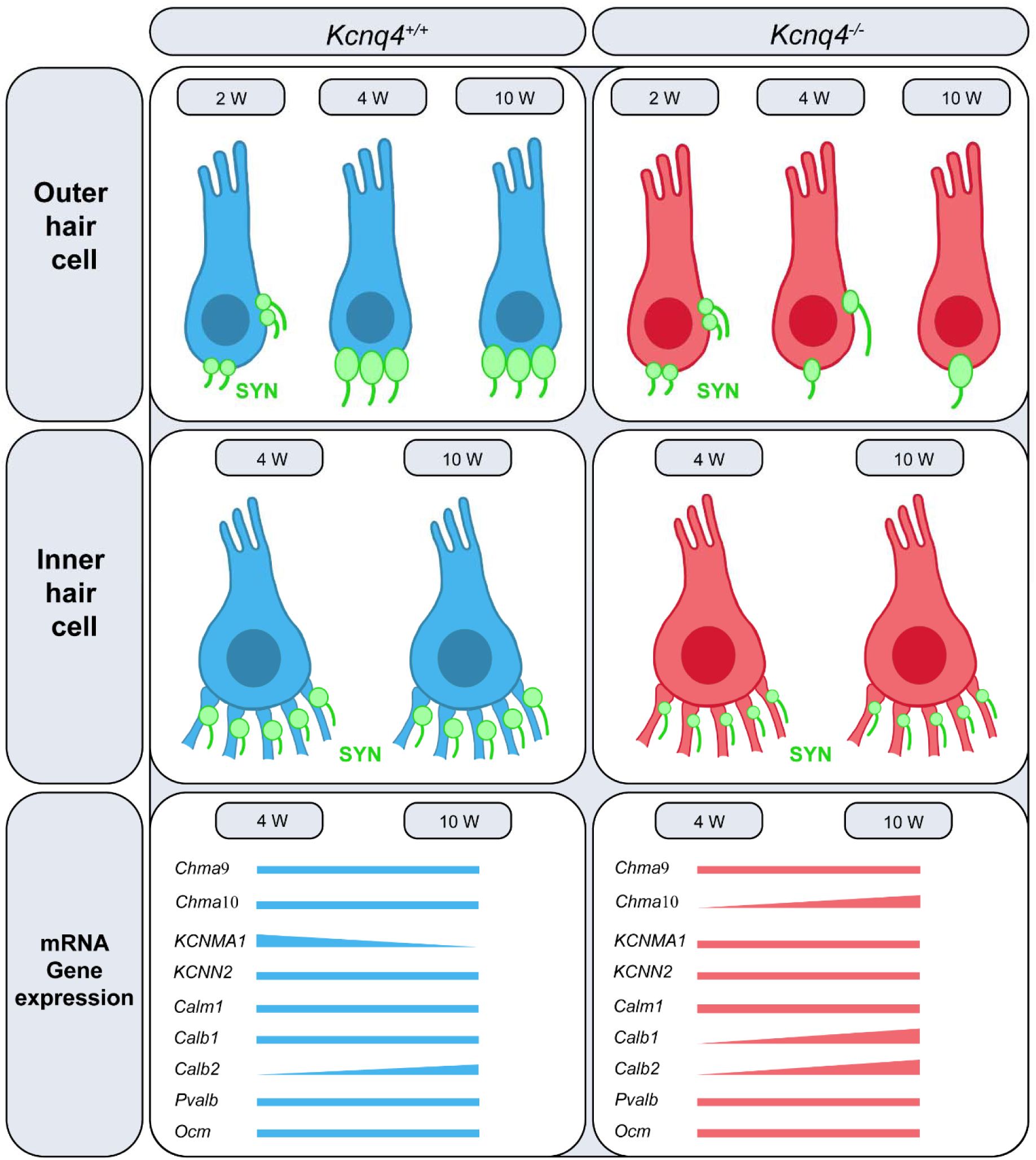

## Introduction

The mammalian auditory system is modulated by two efferent neuronal components descending from the medial and lateral nuclei of the superior olivary complex (MOC and LOC, respectively) [1]. In adults, LOC fibers contact the afferent dendrites that innervate inner hair cells (IHCs), whereas MOC fibers form direct synaptic contacts with outer hair cells (OHCs) [2]. While the precise role of the LOC system remains poorly understood, the function of the MOC system has been extensively characterized. Activation of the MOC pathway inhibits cochlear responses by reducing OHC-mediated amplification, thereby diminishing basilar membrane motility and transiently decreasing auditory nerve sensitivity to sound [3]. In contrast, the LOC system modulates auditory nerve activity directly within the cochlea [4].

During postnatal cochlear development, both IHCs and OHCs undergo extensive remodeling of their afferent and efferent innervation, transitioning from transient, developmentally regulated synaptic contacts to the highly specialized and functionally segregated circuitry characteristic of the mature auditory system [5].

Before the onset of hearing, efferent fibers from the olivocochlear complex invade the cochlear sensory epithelium during late embryonic stages, and at birth efferent synapses are found exclusively around IHCs, while OHCs lack direct efferent innervation [6], [7]. These contacts appear as early as postnatal day 1 in the basal turn of the mouse cochlea [8] and within the first postnatal week in rats [9]. Throughout the first two postnatal weeks, efferent transmission to IHCs is mediated by cholinergic input through the α9α10 nicotinic receptor (nAChR) [10], [11]. Part of this early efferent innervation is transient and undergoes synaptic pruning as maturation proceeds, following basal to apical and modiolar to strial gradients [12].

As maturation progresses, direct efferent contacts onto IHCs are withdrawn and efferent control is progressively relocated onto afferent fibers, primarily via the LOC system [13]. This transition is associated with the loss of nAChR and a reduced responsiveness of SK2 and BK channels to ACh in IHCs [10], [14], [15]. During the second postnatal week, MOC fibers contact OHCs and remain exclusively as direct synaptic inputs to OHCs [11], [16]. MOC fibers exclusively release ACh, generating inhibitory currents [17], although this inhibition is initiated by a transient excitatory current mediated by the α9α10 nAChR [10], [14], [15].

In OHCs, the α9α10 nAChR is a calcium (Ca⁺^2^)- permeable cationic channel [18], [19]. Ca⁺² influx through this receptor activates Ca⁺²-dependent potassium (K⁺) channels (BK and SK2), leading to OHC hyperpolarization via K⁺ efflux [20], [21]. This Ca^+2^ entry is thought to trigger calcium-induced calcium release from nearby subsynaptic cisterns, further promoting SK2 channel activation [22]. In contrast, BK channels exhibit a lower Ca⁺² affinity and therefore require higher intracellular Ca^+2^ concentrations for activation [23], [24]. Notably, the expression of these channels follows a tonotopic gradient along the cochlea: BK channel expression increases from the apical to the basal region, whereas SK2 channels display an opposite gradient [20], [25], [26].

The basal pole of OHCs contains multiple components essential for repolarization and K⁺ handling, including KCNQ4, BK, and SK2, as well as proteins associated with synaptic transmission. Among these, KCNQ4 serves as the primary contributor to OHC repolarization by driving the efflux of excess K⁺ entering during mechanoelectrical transduction [27], [28]. BK and SK2 channels participate in this repolarization process but are specifically engaged during efferent activity via Ca⁺² influx [29].

As mentioned above, calcium influx is a key process in efferent synaptic transduction. The magnitude and spatiotemporal dynamics of Ca⁺² signals in hair cells (HCs) are tightly regulated by calcium-binding proteins (CBPs). The major CBPs in the inner ear (calbindin, calretinin, and parvalbumins) are particularly enriched in HCs and may serve a protective function by buffering intracellular Ca⁺² increases [30]. Calmodulin, another key CBP, indirectly modulates the activity of KCNQ4, BK, and SK2 channels [31]. These mechanisms are critical, as efferent modulation of OHC activity helps prevent excessive depolarization, particularly during sound overstimulation [1]. Supporting this idea, a knock-in mouse model expressing a gain-of-function α9 nAChR subunit showed enhanced efferent tone that confers protection against noise-induced damage and age-related hearing loss [32], [33].

Previous work from our laboratory demonstrated that mice of the C3H/HeJ strain lacking the KCNQ4 channel exhibit progressive degeneration of OHCs, IHCs, and spiral ganglion neurons with age [34]. To elucidate the mechanisms driving this process, we assessed the activation of apoptosis and identified the intrinsic pathway as the primary mediator of HC death [35]. In this model, defective K⁺ extrusion leads to K⁺ accumulation and chronic depolarization of HCs [27]. We hypothesized that this persistent depolarization not only disrupts membrane domains [35] but also efferent synapse organization and alters intracellular Ca⁺² handling, ultimately triggering apoptotic pathways.

To better understand the mechanisms underlying this degenerative process, we investigated the effect of KCNQ4 absence on the localization, number, and volume of efferent terminals in both OHCs and IHCs of the organ of Corti (OC) across immature and mature developmental stages. Additionally, we examined its impact on the expression of several efferent components and proteins involved in calcium homeostasis.

## Methods

### Animal model

C3H/HeJ transgenic mice, lacking the expression of the KCNQ4 protein (*Kcnq4^-/-^*), due to a deletion spanning exon 6 to exon 8, were used [27], [34], [36]. We chose this strain since is relatively resistant to age-related hearing loss [35], [37]–[39]. Wild-type (WT, *Kcnq4^+/+^*) C3H/HeJ litters mice were used as controls. Mice of both sexes were used in all experiments. Animals were divided into age groups: 2, 3, 4, and 10 postnatal weeks (W) for each genotype. The experimental protocol followed in this study was approved by the Council for Care and Use of Experimental Animals (CICUAE, protocol no. 083/2016) of the Universidad Nacional del Sur (UNS), whose requirements are strictly based on the European Parliament and Council of the European Union directives (2010/63/EU).

### Cochlea isolation

Mice were euthanized by CO_2_ exposure, and their inner ears were promptly removed from the temporal bones. Then, were fixed by overnight submersion in 4 % paraformaldehyde (PFA), washed with 1x PBS, and decalcified using 8 % EDTA in 1x PBS for up to 5 days depending on animal age, on a rocking shaker at 4 °C.

### Whole-mount cochlear preparations

The OC from decalcified inner ear was obtained following a procedure previously published [34]. This method consists in splitting the whole cochlear length into three longitudinal segments: basal, middle and apical turns using fine scissors. Then, spiral ligament, modiolus and tectorial membrane were removed and the organ of Corti was isolated. Finally, the hook was cut off from the rest of the basal segment.

### Immunohistochemistry

For immunofluorescence experiments we used middle cochlear turns, since the MOC density innervation to OHCs remains highest in this area [40], [41]. Cochlear turns were postfixed in 4 % PFA during 30 min, washed three times in PBS 1x and incubated for 2 h in blocking solution (2 % BSA, 0.5 % Nonidet P-40 in 1x PBS). Primary antibodies were incubated for 48 h in carrier solution (1x PBS containing 1 % BSA, and 0.25 % Nonidet P-40). Subsequently, tissue was rinsed three times in 1x PBS. Secondary antibodies, diluted in the carrier solution, were incubated for 2 h at room temperature. After that, samples were washed three times in 1x PBS. Finally, cochlear turns were mounted in Fluoromount-G (Southern Biotech). The following primary antibodies were used: mouse anti-synaptophysin (SYN) (1:500; MAB5258, Chemicon), mouse anti-MaxiKα (BK) (1:250; sc-374142, Santa Cruz Biotechnology), mouse anti-calretinin (1:100, sc-365956, Santa Cruz Biotechnology) and mouse anti-calbindin D28K (1:100, sc-365360, Santa Cruz Biotechnology). The following fluorescently-labeled secondary antibodies were used diluted (1:500): donkey anti-rabbit Alexa Fluor 488 (cat#A-21206), donkey anti-mouse Alexa Fluor 488 (cat#A-21202), and donkey anti-mouse Alexa Fluor 555 (cat#A31570). Nuclei were stained with DAPI (1:1000).

### Confocal microscopy and 3D image reconstruction

Confocal z-stacks (∼ 20 µm range size, with 0.5 µm step size) of the medial turn (frequency range ∼ 17- 26 kHz) from each cochlea were taken using a Zeiss LSM900 with an Airyscan module II equipped with 63x oil-immersion lens (1.6x digital zoom for OHC and 1.2x digital zoom for IHC). Each image usually contained 20 to 40 OHCs or 10 to 15 IHCs.

For image visualization shown in the different figures (1, 2, 3 and 4) Regions of Interest (ROIs) of the same height and width were selected for each condition and the final 3D-images were reconstructed with the Surface Projection of the Zeiss ZEN 3.0 program.

### Image processing and efferent terminals analysis

The number of efferent presynaptic puncta, labeled by SYN and post-synaptic puncta, labeled by BK, were counted manually using the Zeiss ZEN 3.0 program, with the Surface Projection of the 3D image reconstruction. The threshold for staining identification was manually set on a per-sample basis at a level at which protein signals of the expected size, shape and cellular location were clearly distinct from the background. For the volume analysis, z-stack images were processed with FIJI Image Analysis software. We measured the total volume of all efferent terminals located within user-defined regions of the cochlea (OHC or IHC region). Next, we used the average value of efferent terminal per HC to calculate indirectly the average individual volume of each puncta.

By using SYN, we interpreted the signal observed in the OHC region as MOC innervation, while the signal in the IHC region was considered to represent LOC innervation, although a small percentage of MOC fibers innervate afferent type I fibers [42], [43].

### Calretinin and calbindin analysis

Z-stacks confocal images labeled by calretinin or calbindin in IHC and OHC from mice of 4 and 10 W were acquired with the configuration mentioned above (same laser and digital gain for each genotype). Images were then analyzed using FIJI Image Analysis software. Maximum intensity projections (MIP) of the individual channels were merged into a 32-bit image which was subsequently thresholded and converted into binary images. The threshold for staining identification was the same between conditions and manually selected at a level at which protein signals of the expected size and shape were clearly distinct from the background. The Mean Fluorescense Intensity (MFI) of each cell was measured by the FIJI software, and the MFI from 10- 20 cells from each cochlear sample was averaged and expressed as arbitrary units (a.u.) (n = 5 for each genotype).

### RNA isolation and Reverse Transcription

Cochlear RNA was extracted from 4 and 10 W mice according to Rías *et al*. (2025) [35]. For each experiment, samples from three mice were pooled. In brief, immediately after cochlear excision, tissue was immersed in an RNA preservation buffer and then, the vestibular region was removed to keep only the cochlea. Total RNA was isolated using the Bio-Zol reagent (PB-L Productos Bio-Lógicos, RA0202) in combination with chloroform and isopropanol protocol. cDNA was produced from 500 ng total RNA with MMLV Transcripta (PB-L Productos Bio-Lógicos, EA1301) using anchored oligo (dT)s following manufacturer’s indications.

### Quantitative PCR

Quantitative PCR (qPCR) was carried out using the cDNA generated previously employing the SsoAdvanced Universal SYBR Green Supermix (BioRad, cat#1725271) in a Rotor-Gene 6000 real-time PCR cycler (Quiagen). Table 1, shows the list of primers used for qPCR detection of the detailed gene expression. As reference genes, we used *GAPDH* and *HPRT*. Messenger RNA (mRNA) subunit expression was referred to the geometric mean of *GAPDH* and *HPRT*. Data analysis was done applying the ΔΔCt method [44], [45] to obtain the relative mRNA quantification (RQ).

**Table 1.**
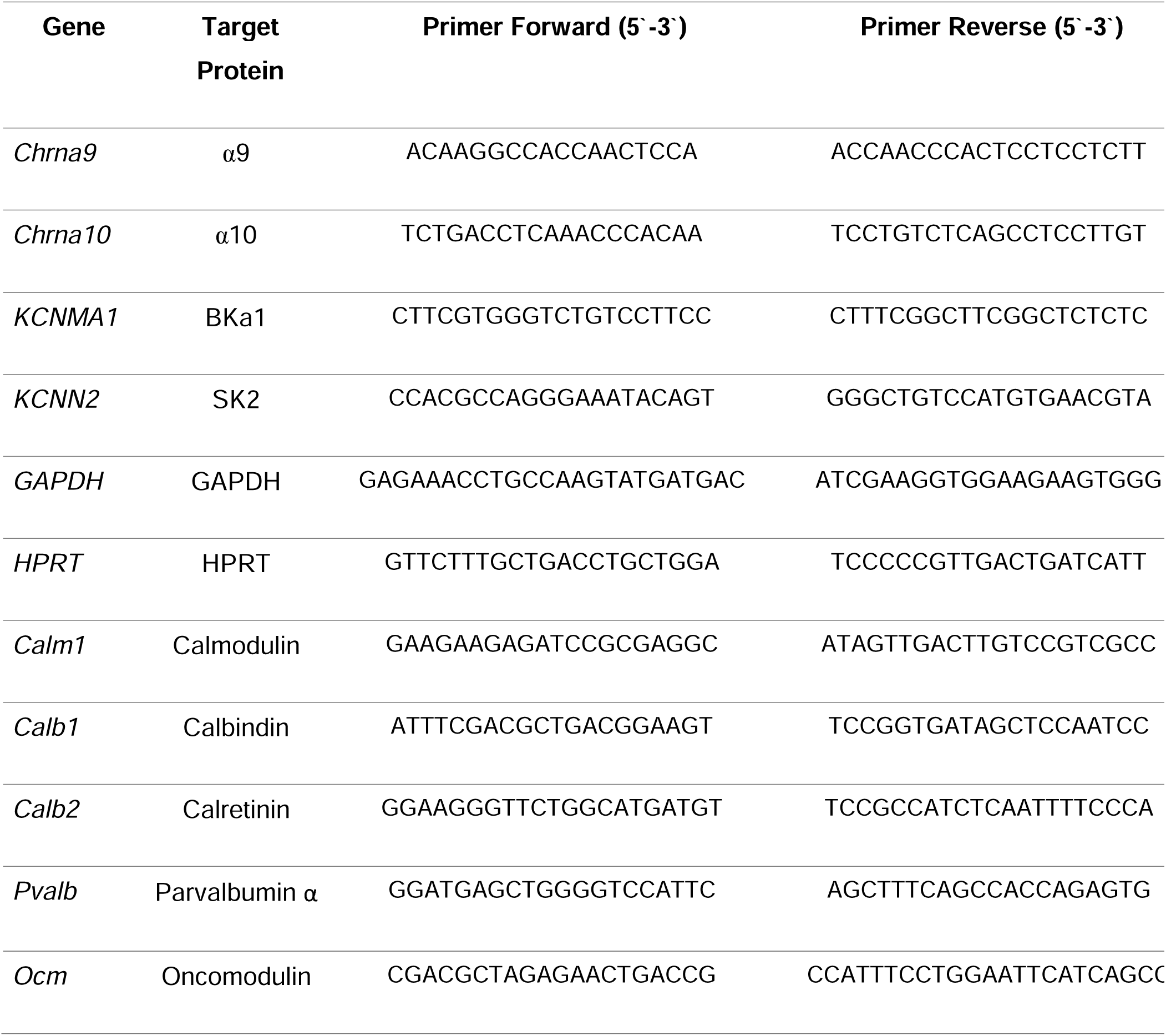
List of primers used for qPCR experiments.

### Western blot

Cochlear proteins were extracted at 4 and 10 W mice for both genotypes. For each condition, cochleae from two mice were pooled. Cochleae were promptly dissected and placed in a 1.5 mL eppendorf tube with 100 μL lysis buffer (5 mL 1M Tris pH 7.4, 5 mL 1M NaCl, 0.5 mL 0.5M EDTA, 2.5 mL 100X triton, 237 mL miliQ H_2_0) with protease inhibitor cocktail (Roche, cat. #05892791001) and homogenized with a pestle (JBF-CSP001002) on ice. Samples were the centrifuged at 12,000 rpm for 10 minutes at 4 °C (Yingtai Instrument TGL16E). The supernatant was extracted and placed into a new eppendorf tube and centrifuged again for 10 minutes at 4 °C. After centrifugation, protein was quantified by Bradford method and 80 µg of protein was resolved by 10 % SDS-polyacrylamide gel electrophoresis and transferred onto a PVDF membrane (Osmonics Inc, Gloucester, MA). The membrane was blocked and incubated with specific antibodies to anti-MaxiKα (1:500; sc-374142, Santa Cruz Biotechnology) and anti-GAPDH (1:1000; sc-32233, Santa Cruz Biotechnology) at 4 °C overnight followed by incubation with HRP conjugated secondary antibody for 1 h at room temperature. The chemiluminescent reaction was developed using the ECL detection system and then the bands were revealed in the high-quality chemi imaging and gel documentation system GBox Chemi XRQ with the G-plot (Syngene). Western blot bands relative density was calculated with Image J free software and expressed in relative units (RU) (https://imagej.nih.gov/ij/).

### Statistics analysis

All data and results were confirmed through three or more independent experiments (indicated for each figure). Values are presented as means ± SEM. Significant differences were identified using Student’s t-test (Fig. 1, 4, 5 and 6) and one- or two-way ANOVA (Fig. 2 and 3) depending on the type of comparison. Tukey’s Multiple post hoc tests were used when ANOVA detected differences. Statistical analyses were performed using Graph Pad 9.0 Software. Outlier values were excluded when detected (Alpha = 0.0500). Statistical significance is represented with (*) when p < 0.0500; (**) when p < 0.0100; (***) when p < 0.0010 and (****) when p < 0.0001.

**Fig. 1.**
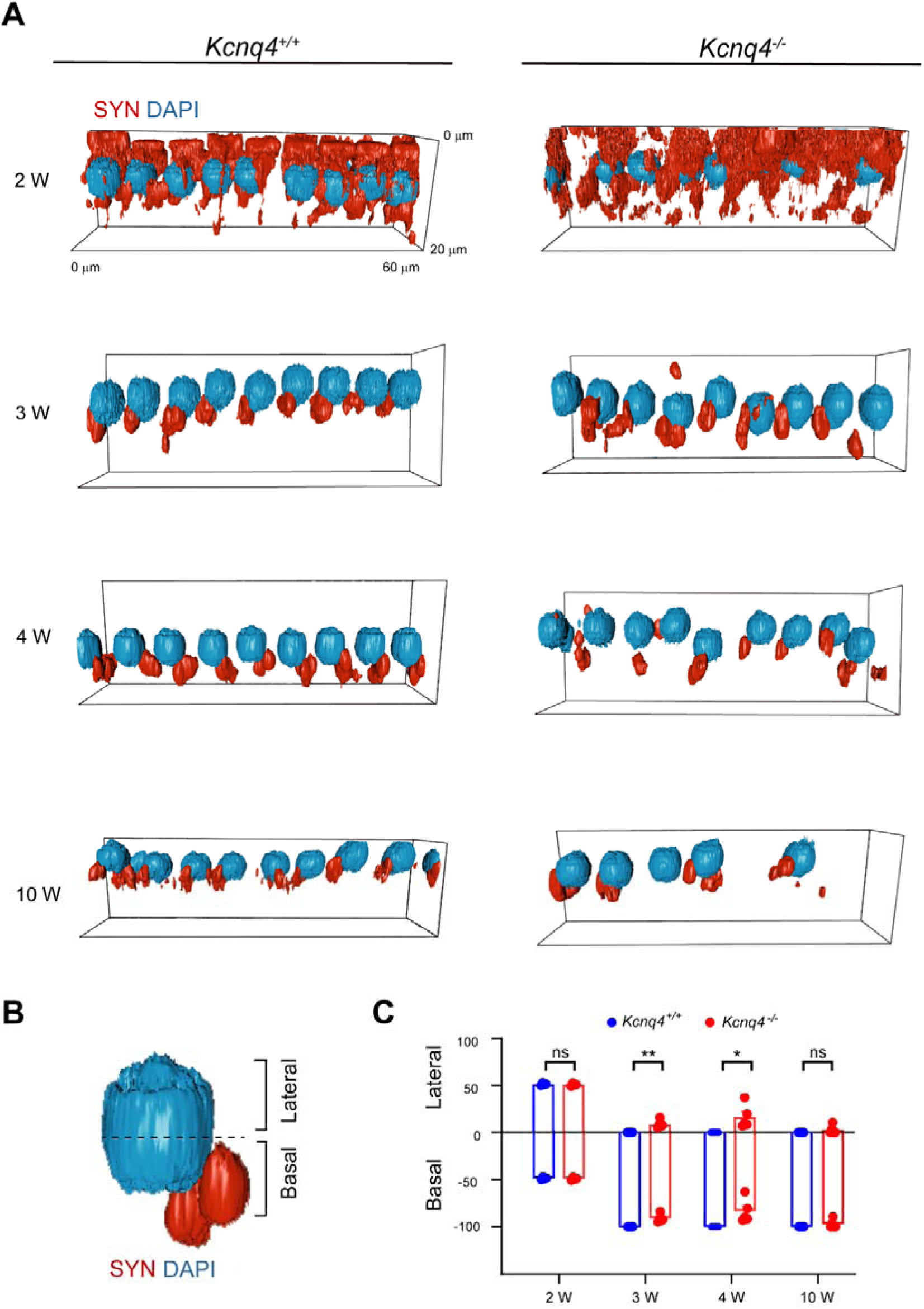
Spatial distribution of efferent terminals in OHCs of KCNQ4 KO mice. **A.** Confocal images from whole-mount mice OC, showing synaptophysin (SYN, red) and nuclei (DAPI, blue) of OHC inner row in the middle cochlear turn for both genotypes at different ages. **B.** Magnification of a single OHC, depicting the basal and lateral domains. **C.** Quantification of the number of total MOC terminals in each domain expressed as a percentage. Data are presented as mean ± SEM. Two-way ANOVA and Tukey’s post hoc test were used. ns, not significant. *, p < 0.0500. **, p < 0.0100.

**Fig. 2.**
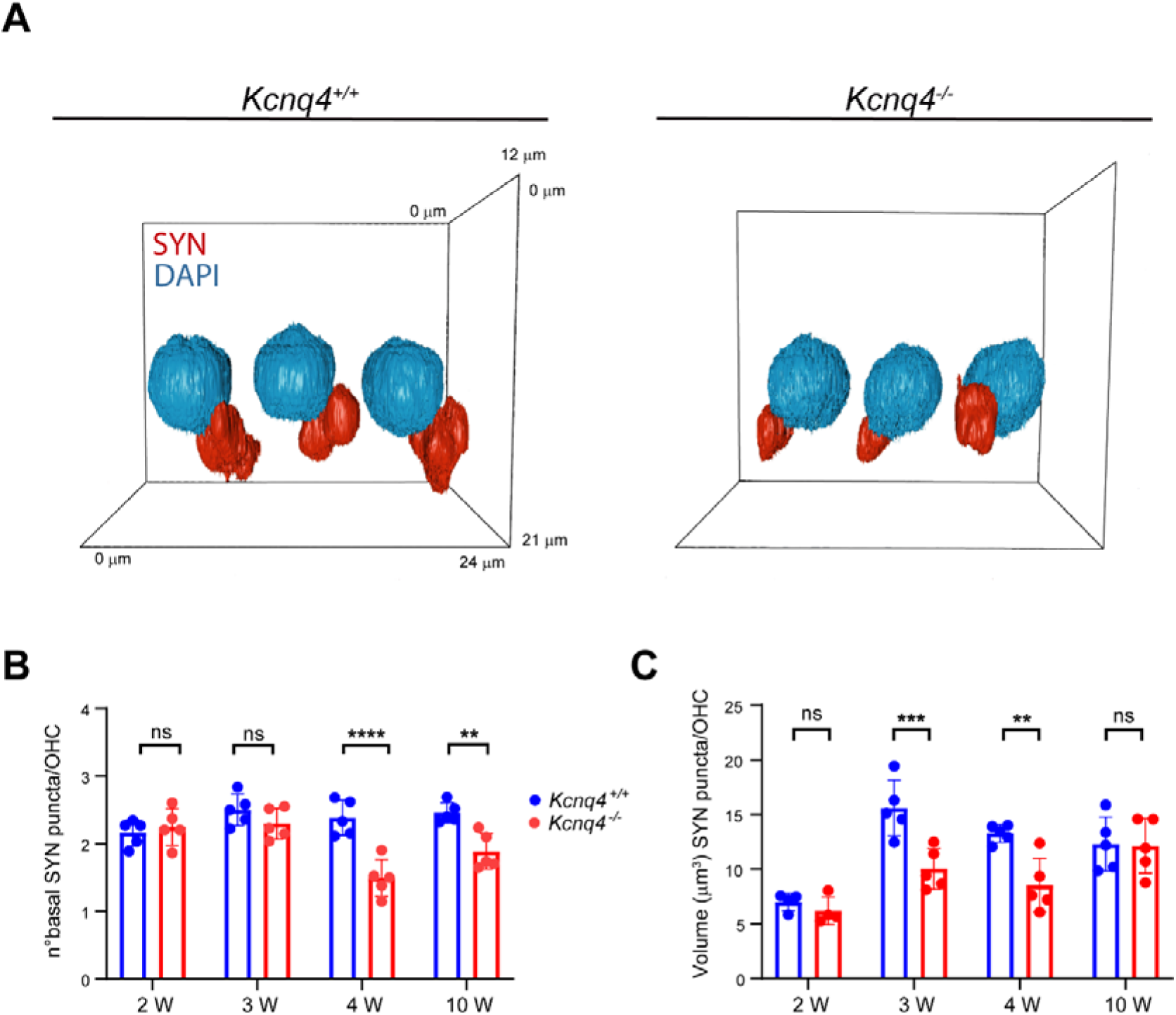
Morphometric analysis of efferent terminals in OHCs of KCNQ4 KO animals. **A.** Magnification of confocal images from a whole-mount OC showing synaptophysin (SYN, red) and nuclei (DAPI, blue) of OHC in the inner row of middle cochlear turn for both genotypes at 4 W. Scale bar set on image. **B.** Quantification of the number of SYN puncta per OHC in the basal domain for WT and KO animals at different ages. Data are presented as mean ± SEM. Two-way ANOVA and Tukey’s post hoc test were used. ns, not significant. **, p < 0.0100. ****, p < 0.0001. **C.** Quantification of the volume of each SYN pucta in OHC for WT and KO animals at different ages. Data are presented as mean ± SEM. Two-way ANOVA and Tukey’s post hoc test was used. ns, not significant. **, p < 0.0100. ***, p < 0.0010.

**Fig. 3.**
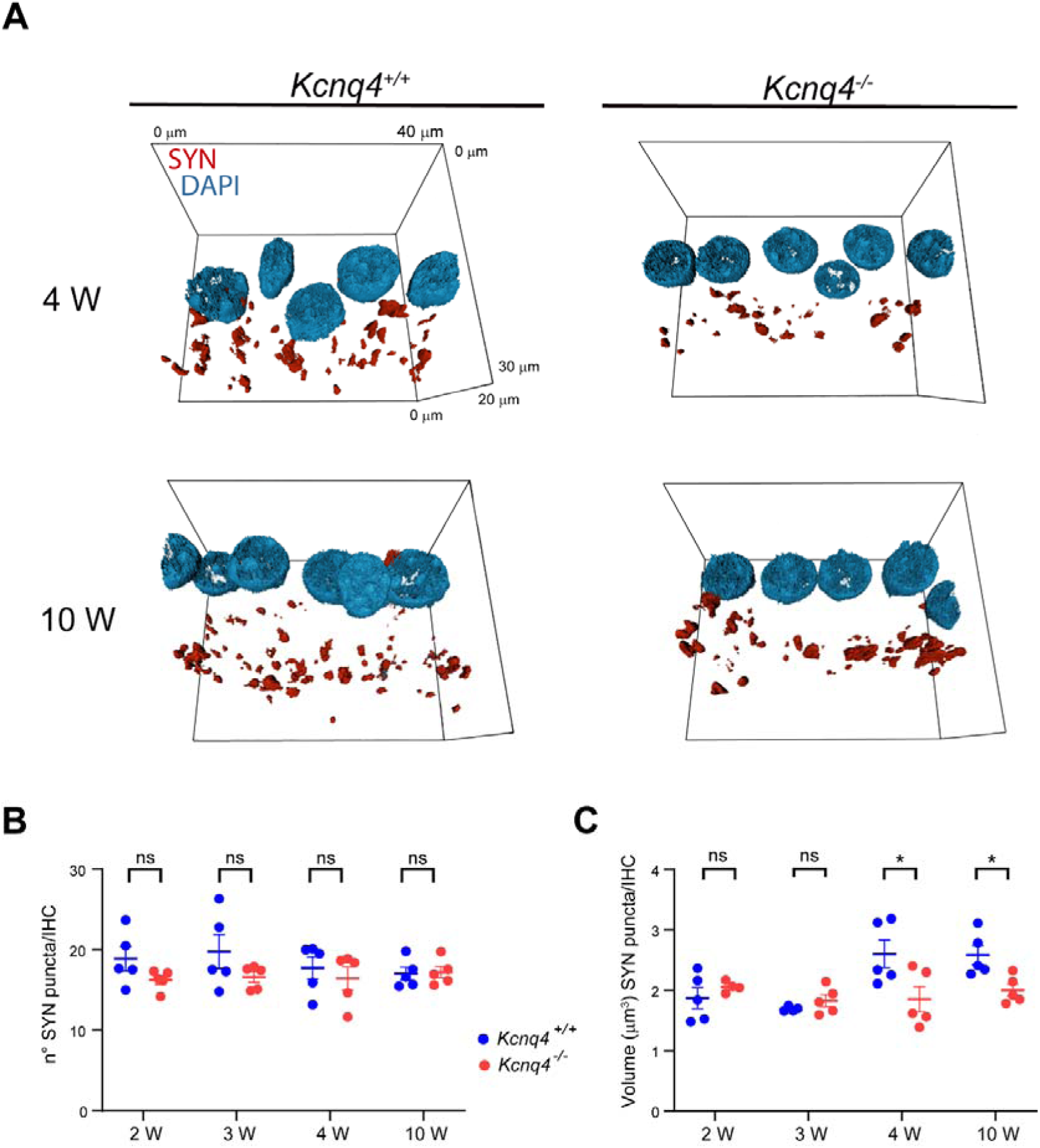
Analysis of efferent terminals in IHCs area in *Kcnq4^-/-^* mice. **A.** Magnification of confocal images showing synaptophysin (red) and nuclei (blue) of IHC in the middle turn for both genotypes at 4 W and 10 W. Scale bar set on image. **B.** Quantification of the n° of SYN puncta in the IHC region, for WT and KO animals at different ages. Data are presented as mean ± SEM. Two-way ANOVA was used. ns, not significant. **C.** Quantification of the volume of SYN puncta in IHC for WT and KO animals at different ages. Data are presented as mean ± SEM. Two-way ANOVA and Tukey’s post hoc test were used. ns, not significant. *, p < 0.0500.

**Fig. 4.**
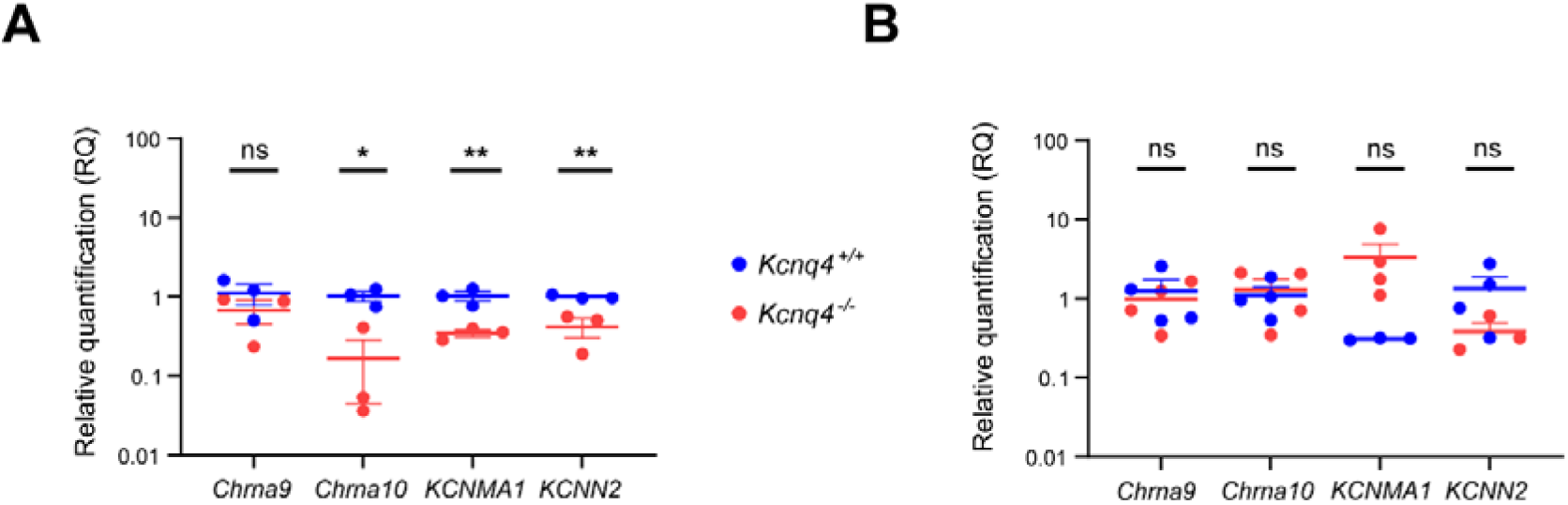
Gene expression of HCs’ efferent components in KCNQ4 KO mice. Relative quantification of gene expression in the OC of α9 (*Chrna9*) and α10 (*Chrna10*) nAChR subunits, BK (*KCNMA1*), and SK2 (*KCNN2*) mRNA in WT and KO animals at 4 W (**A**) and 10 W (**B**). Data are expressed as the mean ± SEM. *GAPDH* and *HPRT* were used as housekeeping genes. Student’s t-test was applied for each gene. ns, not significant. *, p < 0.0500. **, p < 0.0100.

**Fig. 5.**
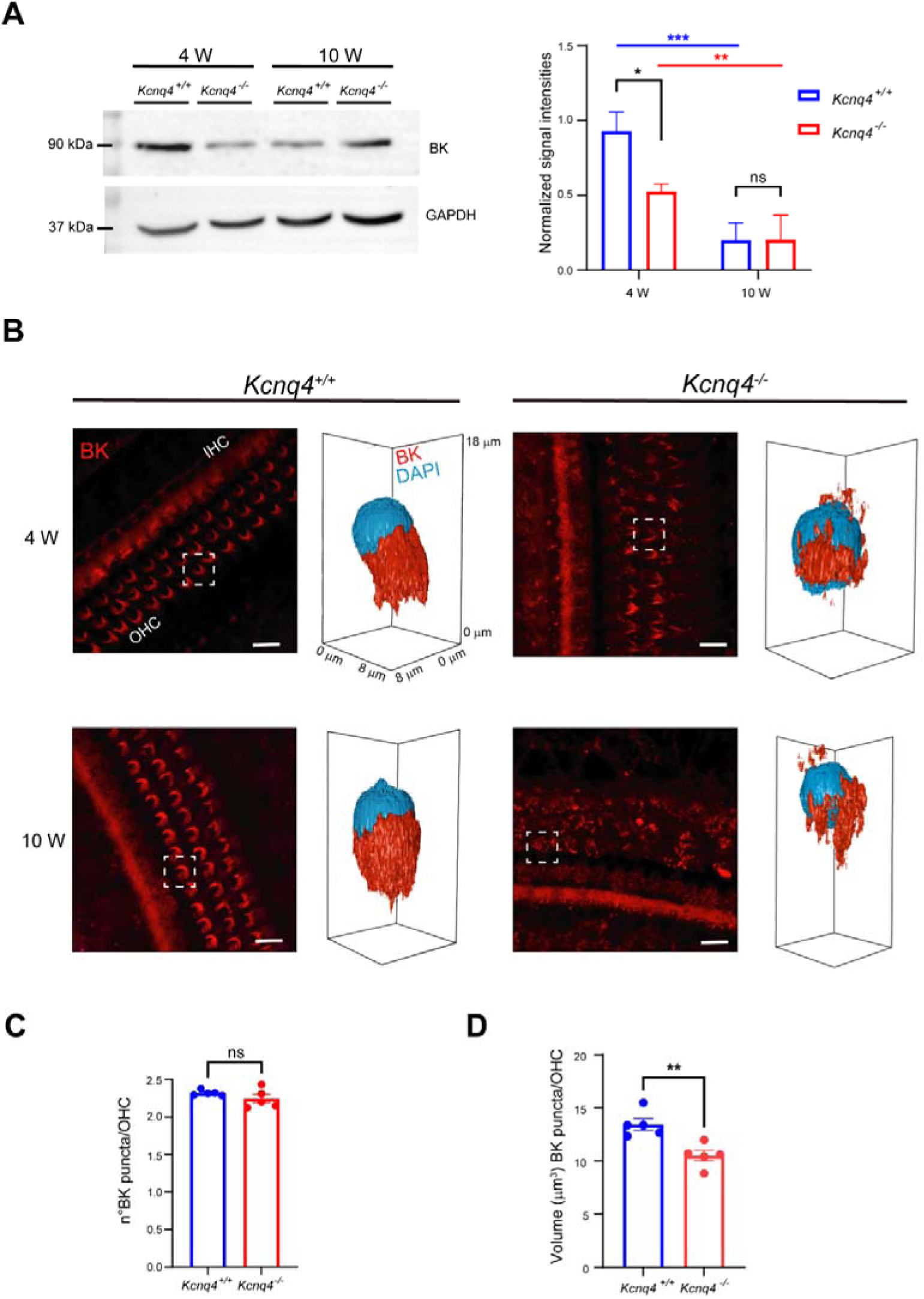
BK expression in the OC of KCNQ4 KO mice. **A.** Representative Western blot analysis of BK (90 kDa band) and GAPDH (37 kDa band) from WT and KO at 4 W and 10 W (left). Quantification of BK band intensity normalized by GAPDH (right). Data are presented as mean ± SEM. Unpaired t-test was used. ns, not significant. *, p < 0.0500. **, p < 0.0100. ***, p < 0.0010. **B.** Confocal images showing BK (red) signal in OHC in the middle turn for both genotypes at 4 and 10 W. Scale bar: 10 μm. Inset depicting the surface confocal image of BK (red) and DAPI (blue) in a single OHC. Scale set on image. **C.** Quantification of the number of BK puncta per OHC in the basal domain for WT and KO animals at 4 W. Data are presented as mean ± SEM. Unpaired t-test was used. ns, not significant. **D.** Quantification of BK puncta volume per OHC in the basal domain for WT and KO animals at 4 W. Data are presented as mean ± SEM. Unpaired t-test was used. **, p < 0.0100.

## Results

### Spatial location of efferent terminals in OHC

During the maturation process, efferent synaptic contacts, initially found in the different membrane compartments (basal, lateral and apical domains) of the OHC, reorganized to the basal domain, around 3 W [46]. To gain insight into the spatial localization of efferent terminals along OHCs in the chronically-depolarized OHCs of the KO mice, we performed cochlear immunofluorescence analysis of SYN, a key component of the synaptic vesicles located in efferent presynaptic terminals within the OC [47]. Using confocal imaging, we evaluated the localization of MOC terminals on OHC in *Kcnq4*^+/+^ and *Kcnq4*^-/-^ animals at different developmental stages: 2 W (hearing onset) [5], 3 W (mature OHC innervation) [46], 4 W (tissue maturation) [48], and 10 W (auditory system fully developed) [49] (Fig. 1A). Three dimensional reconstructions of SYN and DAPI signals revealed the characteristic rounded morphology of efferent synaptic boutons contacting OHCs (Fig. 1A). To quantify the number of efferent boutons in each domain we defined the basal domain as the region located below the nucleus equator and the lateral domain as the region above this imaginary line (Fig. 1B). Next, we counted the number of MOC terminals located between each domain at the different ages analyzed. At 2 W, both genotypes possessed ∼ 51 % of synaptic boutons in the lateral domain and ∼ 49 % in the basal domain (Fig.1C). Subsequently, from 3 W onwards, all terminals relocated to the basal domain in WT animals, consistent with the fact that MOC terminals are exclusively located in the basal domain of OHC at mature stages [5], [46] (Fig. 1C, blue bars). In contrast, in the KO OHCs this shift is significatively altered at 3 and 4W, with 8.7 % and 16.5 % of the terminals persisting in the lateral domain at 3 and 4 W, respectively (Fig. 1C, red bars). By 10 W, in *Kcnq4*^-/-^ animals, innervation appears to reach mature values (p = 0.0450, One-way ANOVA, Tukey’s post hoc test), compared to 3 and 4 W. These observations indicate that the normal development of the efferent terminals may be delayed in *Kcnq4*^-/-^ animals.

### Morphometric characteristics of efferent terminals in OHC

We evaluated the number and volume of the synaptic boutons in the basal domain of OHCs, which corresponds to the region where synapses are normally located following maturation [46]. We observed a typical rounded morphology for single terminals for both genotypes (Fig. 2A). In WT mice, the number of presynaptic contacts per OHC at all ages remains constant and was ∼ 2.2 (p = 0.0965, One-way ANOVA) (Fig. 2B, blue bars). On the contrary, for KO animals, the number of presynaptic boutons changed with age (p = 0.0005, One-way ANOVA). Comparing WT and KO mice at each age, the number of presynaptic contacts per OHC was similar to their corresponding controls at 2 W and 3 W (p = 0.9839 and p = 0.4984, respectively, Two-way ANOVA, Tuckey post hoc test), but decreased to less than 2 puncta/ OHC in KO mice at 4 and 10 W (p < 0 .0001 and p = 0.0017, respectively, Two-way ANOVA, Tuckey post hoc test) (Fig. 2B, red bars).

In addition, we studied if the volume of the efferent presynaptic puncta would also be affected with maturation. We found that the volume per bouton at the immature stage of 2 W was similar for both genotypes (approximately 7 μm^3^, p = 0.9984, Two-way ANOVA, Fig. 2C). At more mature stages, presynaptic terminals from WT animals fastly duplicate their size at 3 W (p < 0.0001, One-way ANOVA, pos hoc Tukey test), which remained stable thereafter (p = 0.2672, One-way ANOVA) (Fig. 2C, blue bars). In contrast, KO mice showed no significant change in the puncta volume during maturation (p = 0.0552, One-way ANOVA) until 10 W, when a volume comparable to that of WT animals was reached (Fig. 2C, red bars). However, there were differences between genotypes at 3 and 4 W (p = 0.0003 and p = 0.0010, respectively, Two-way ANOVA, Fig. 2C).

These results suggest that the absence of KCNQ4 in OHCs leads to transient alterations in the spatial localization, number and volume of the efferent synaptic terminals, which could imply a delayed maturation process.

### Morphometric characteristics of efferent boutons in the IHC region

In addition to the analysis of MOC efferent terminals in OHCs, we performed the same analysis in the LOC efferent fibers that innervate the afferent fibers beneath the IHCs. Although a small percentage of MOC fibers innervate afferent type I fibers [42], we considered SYN signal in the IHC region as LOC innervation [43]. We analyzed the number of the presumed LOC efferent synaptic boutons across different ages. In both genotypes, we observed a greater number of smaller efferent boutons beneath IHCs compared to those seen in OHCs (Fig. 3A). At 2 W, the number of efferent boutons per IHC was ∼ 19 in WT animals and ∼ 16 in KO animals, with no significant difference observed (p = 0.8021, Two-way ANOVA; Fig. 3B). These values remained consistent up to 10 W for both genotypes (p= 0.5998 for WT and p = 0.8905 for KO, One-way ANOVA for each) (Fig. 3B).

Similarly, the volume of each synaptic bouton was evaluated. At 2 W, both genotypes exhibited volumes of ∼ 1.9 μm^3^, with no significant differences between them (p= 0.8888; mixed effects model, Fig. 3B). At 3 W, bouton volumes were around 1.7 μm^3^ for both groups, again with no significant differences (p = 0.9584; mixed effects model, Fig. 3B). The comparison between 2 and 3 W within each genotype revealed no significant differences (p = 0.8540 for WT and p = 0.7555 for KO, respectively; Two-way ANOVA, Fig. 3B).

At 4 W, WT mice showed a significant increase in average bouton volume (∼ 2.6 μm^3^) compared with younger ages (p = 0.0099 and p = 0.0020, respectively; Two-way ANOVA, Tukey’s post hoc test, Fig. 3C), and this value remained stable up to 10W (p = 0.9998; Two-way ANOVA, Fig. 3C). In contrast, 4 W KO animals, maintained bouton volumes similar to those observed at 2 and 3 W (∼ 1.8 μm^3^) (p = 0.8105 and p = 0.9995, respectively, Two-way ANOVA), with no changes up to 10 W (p= 0.8957; Two-way ANOVA, Fig. 3C). When comparing average bouton volumes between genotypes, significant differences were detected at 4 W (p = 0.0060; mixed effects model, Sidak’s post hoc test, Fig. 3C) and also at 10 W (p = 0.0435; mixed effects model, Sidak’s post hoc test, Fig. 3C).

These results suggest that LOC efferent terminals in KO animals also suffer a mild morphometric alteration, as evidenced by a slightly reduced presynaptic bouton volume compared to WT mice.

### Expression of synaptic efferent components

Our results indicate that efferent presynaptic boutons exhibit altered morphology and location in the absence of KCNQ4. To gain insight whether this absence also affects the expression of efferent postsynaptic components, namely channels and receptors, we performed qPCR on inner ear samples from 4 and 10 W animals of both genotypes. We evaluated the relative mRNA expression of the potassium channels BK and SK2, as well as the α9 and α10 subunits of the α9α10 nAChR complex, which are mainly expressed at the efferent postsynaptic membrane [20]. We found a significant decrease in the gene expression level of the α10 nAChR subunit at 4 W in KO animals (p = 0.0107, Student’s t-test, Fig. 4A), returning to WT levels at 10 W (p = 0.7172, Student’s t-test, Fig. 4B). For α9 nAChR subunit gene, no significant expression changes were detected at any age in either genotype (p = 0.3342 and p = 0.6660, respectively, Student’s t-test) (Fig. 4A and B). Regarding potassium channels, we found a significant reduction in BK and SK2 mRNA levels at 4 W in KO animals (p = 0.0099 and p = 0.0082, respectively, Student’s t-test, Fig. 4A), reaching to WT levels at 10 W (p = 0.1403 and p = 0.1972, respectively, Student’s t-test, Fig. 4B). When comparing mRNA levels between 4 W and 10 W for each genotype, a significant difference was observed only for *KCNMA1* (BK) expression in WT animals, which was reduced by approximately 3.3-fold at 10 W (p = 0.0074, Student’s t-test, Fig. 4).

### BK protein expression in OHC

To better understand the changes in the efferent components of the postsynaptic membrane, we determined BK protein expression in the OC by western blotting for both genotypes at 4 and 10 W (Fig. 5A, left), considering that BK mRNA levels were the only ones showing time-dependent changes (see above). At 4 W, we found a significant decrease in BK protein levels in KO mice compared to WT animals (p = 0.0389, Student’s t-test, Fig 5A, right). Meanwhile, at 10 W, BK protein levels were markedly reduced in both genotypes (p = 0.0098 and p = 0.0098, respectively, Two-way ANOVA), showing no significant differences between them (p = 0.9859, Student’s t-test) (Fig. 5A, right). This reduction was approximately 4.5-fold in WT animals, whereas in KO it was ∼ 2.5-fold.

To further investigate the subcellular organization of BK, we additionally performed whole-mount immunofluorescence analysis of BK at 4 and 10 W. By confocal imaging, in WT mice we observed a characteristic “C shaped” pattern in a top view of each OHC (Fig. 5B) [20], [21], [26], [50]. In the 3D reconstructions, this signal formed plate-like structures composed of two-lobes, as previously described[35] (Fig. 5B). In KO animals, this organization was altered, showing a different spatial distribution that extended over both the lateral and basal domains of the OHC and formed irregular shapes, with a loss of the characteristic plaque-like organization that became more pronounced at 10 W (Fig. 5B, right).

Based on these observations, we then analyzed the number of BK puncta per OHC. At 4 W, we quantified an average of ∼ 2.32 BK puncta/OHC in both genotypes (p = 0.2399, Student’s t-test, Fig. 5C). At 10 W in the WT animals, we observed a slight but significant decrease in the number of puncta per OHC (mean = 2.19, p = 0.0461, Student’s t-test). Due to the highly irregular expression pattern observed at 10 W in KO animals, BK puncta could not be reliably quantified.

We also determined BK puncta volume. At 4 W, WT animals presented a mean volume of ∼ 13.50 μm^3^ per OHC, which was significantly larger than the ∼ 10.50 μm^3^ observed in KO mice (p = 0.0042, Student’s t-test, Fig. 5D). At 10 W for WT animals, BK signal volume showed no differences compared to 4 W (mean = 15.23, p = 0.2483, Student’s t-test). As noted above, BK volume per OHC in KO animals could not be quantified at this age.

In summary, gene expression and protein levels for BK channels changes with maturation in both genotypes, but there were clear differences in the expression levels between WT and KO and the localization and organization of the channel in the OHC membrane.

### Calcium binding proteins analysis

As previously described, Ca^+2^ influx in the HCs is regulated by different CBPs [30]. To further investigate CBPs expression in HCs, we performed qPCR studies in 4 W and 10 W animals. The genes encoding some of the main proteins involved in this process were analyzed: *Calm1* (calmodulin) (ubiquitous), *Calb 1* (calbindin) (both HCs, and spiral ganglion neurons) [51], *Calb 2* (calretinin) (mainly IHC expression), *Pvalb* (parvalbumin α) (OHC expression), and *Ocm* (oncomodulin) (OHC expression). At 4 W, no significant differences in mRNA expression levels were detected between genotypes for any of the genes examined (p > 0.0500 for each one, Student’s t-test) (Fig. 6A). In contrast, at 10 W, *Calb1* and *Calb2* mRNA levels were significantly elevated in *Kcnq4^-/-^* animals compared to WT, with increases of approximately 10-fold and 20-fold, respectively (p = 0.0321 and p = 0.0047, respectively, Student’s t-test, Fig. 6B).

**Fig. 6.**
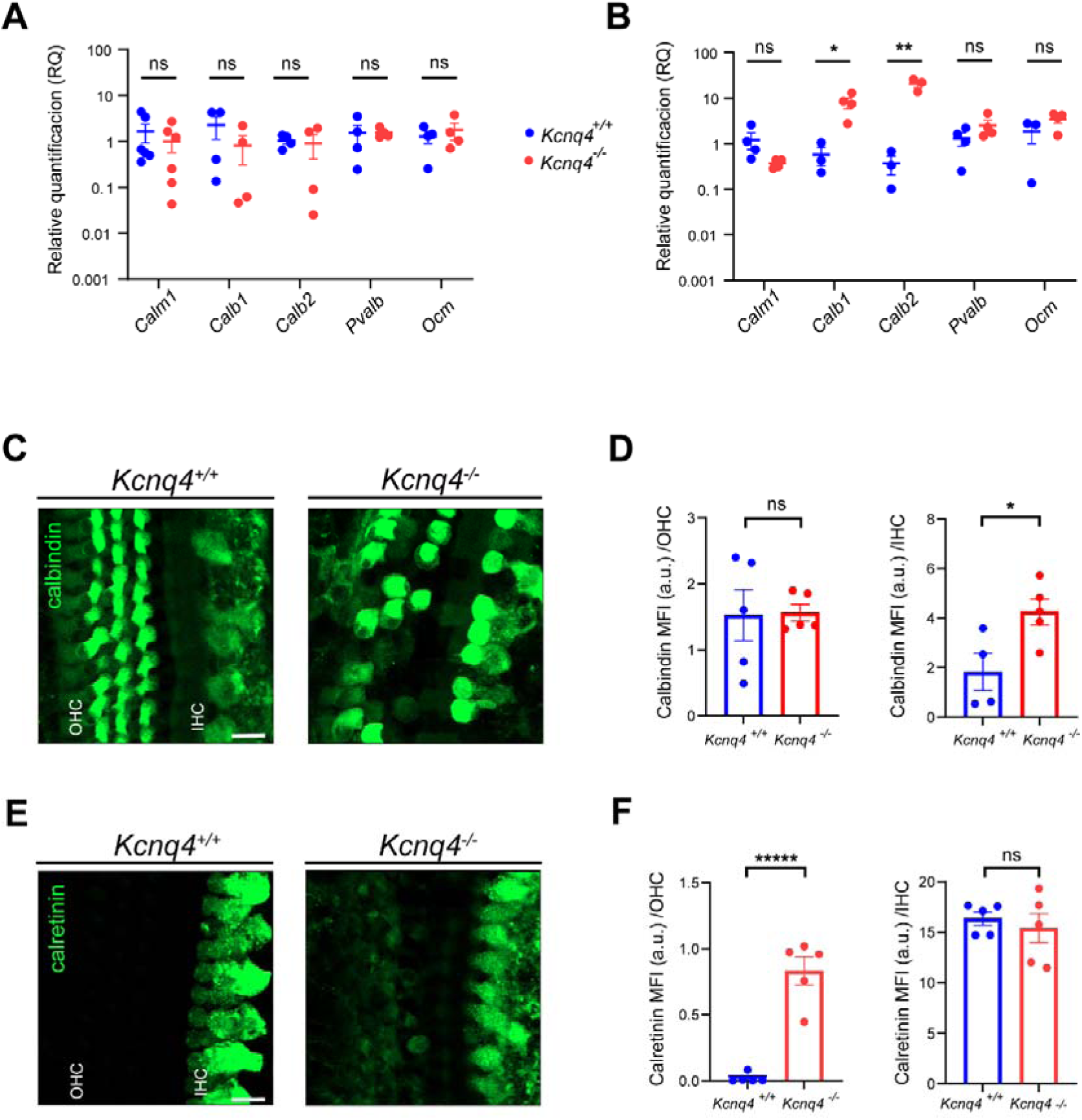
Calcium binding proteins analysis in the OC of mice lacking KCNQ4 expression. **A,B.** Relative quantification of calmodulin (*Calm1*), calbindin (*Calb1*), calretinin (*Calb2*), parvalbumin α (*Pvalb*) and oncomodulin (*Ocm*) mRNA in the OC of WT and KO animals at 4 W (**A**) and 10 W (**B**). Data are expressed as the mean ± SEM. GAPDH and HPRT were used as housekeeping genes. Student’s t-test was applied for each gene. ns, not significant. *, p < 0.0500. **, p < 0.0100. **C.** Confocal images showing calbindin (green) signal in IHC and OHC, in the middle cochlear turn for both genotypes at 10 W. Scale bar: 10 μm. **D.** Quantification of calbindin levels expressed as mean fluorescence intensity (MFI) in OHC and IHC, respectively. Data are presented as mean ± SEM. Student’s t-test was used. ns, not significant. *, p < 0.0500. **E.** Confocal images showing calretinin (green) signal in IHC and OHC in the middle cochlear turn for both genotypes at 10 W. Scale bar: 10 μm. **F.** Quantification of calretinin levels expressed as MFI intensity in OHC and IHC, respectively. Data are presented as mean ± SEM. Student’s t-test was used. ns, not significant. ****, p < 0.0001.

To determine whether these transcriptional changes were reflected at the protein level in the cells of the OC, we evaluated the immunofluorescence signal intensity of proteins that showed genotypic differences in mRNA levels at 10 W: calretinin and calbindin. In whole-mount preparations of the OC, we evaluated the expression of these CBPs in both OHC and IHC (Fig 6C and 6E). A strong calbindin signal was observed in both IHCs and OHCs of both genotypes; however, in IHCs, signal intensity was lower in WT animals (Fig. 6C). For calretinin, a predominant signal was detected in the IHC region in both genotypes, whereas in OHCs only a weak signal was observed in KO animals (Fig. 6E).

To quantify the degree of change in protein expression, we determined the mean fluorescence intensity (MFI) for each protein signal. MFI measurements revealed that calbindin exhibited a uniform distribution in both HCs types (Fig. 6D). In OHCs, no genotypic differences were detected (p = 0.9310, Student’s t-test, Fig. 6D left). In contrast, IHC of KO animals exhibited a significant increase in calbindin expression relative to WT (p = 0.0283, Student’s t-test, Fig. 6D right). Quantification of calretinin MFI per cell type revealed that for both genotypes, expression was higher in IHC than in OHC (Fig. 6F). However, in KO mice, OHC displayed a significant increase in calretinin intensity compared to WT (p < 0.0001, Student’s t-test, Fig. 6F left), whereas no differences were observed in IHC (p = 0.5684, Student’s t-test, Fig. 6F right).

Overall, these findings suggest that hair cells undergo adaptive responses to alterations in calcium levels. Genotype-dependent changes in the gene expression of two CBPs in the OC show differential correlations with protein expression levels in IHCs and OHCs.

## Discussion

Considering that sensory HCs are among the few cell types that depolarize due to K^+^ influx at the apical surface exposed to the endolymph, the KCNQ4 channel is essential for hearing, serving as the major pathway for K^+^ efflux from OHCs. Dysfunction of this channel, caused by dominant-negative mutations in the KCNQ4 gene, underlies DFNA2, a human deafness that worsens over years [52], [53]. The pathophysiological mechanism, elucidated through mouse models, involves loss of KCNQ4 function that causes intracellular potassium accumulation due to the endocochlear potential and sustained depolarization, resulting in chronic cellular stress and OHC degeneration [27], [34], [35]. Postnatal maturation of the mouse inner ear involves the expression and dynamic changes in the cellular localization of several proteins that are essential for HC-specific functions during the first 4-6 W after birth [46]. KCNQ4 expression begins around postnatal day 6, initially localizing to the basolateral membrane domain and subsequently redistributing to the basal domain by postnatal day 12 [54], which coincides with the emergence of the characteristic electrophysiological properties of *I_K,n_* current mediated by KCNQ4 [55]. Similarly, prestin under-goes postnatal relocalization to the lateral membrane domain [46]. Notably, the absence of KCNQ4 leads to alterations in the expression pattern of essential proteins, as well as to structural abnormalities in the hair bundles [35]. Our results underscore the critical function of KCNQ4 in safeguarding auditory integrity and efferent innervation. Furthermore, it elucidates the novel significance of KCNQ4 for cochlear development, revealing that its dysregulation contributes to tissue alterations.

### Efferent synapses in OHC

In this work, we found that the efferent synapses undergo age-dependent structural and organizational changes in the absence of KCNQ4. The localization, number, and volume of efferent puncta in OHC change over time in *Kcnq4^-/-^* animals. At the onset of hearing (2 W), the percentage distribution of efferent puncta is similar between both genotypes (WT and KO), with presynaptic terminals located in both the lateral and basal domains. However, as hearing matures, *Kcnq4^-/-^* animals fail to efficiently relocate their MOC terminals to the basal domain, with some terminals persisting in the lateral domain until 10 W. Within this developmental window, our model exhibits impaired hearing across all cochlear regions, as well as degeneration and loss of OHCs [34]. The alterations in the proper localization of efferent terminals observed in the KO animals were analyzed in the middle cochlear turn, where cell loss is minimal, and are likely to contribute to the progressive degeneration of these cells. This mislocalization of MOC terminals in OHC was also observed by Takahashi et al. in adult prestin-KO mice [46]. The authors showed that efferent terminals were present in both the lateral and basal domains at both immature and mature stages in mice lacking prestin [46]. Similarly, in CaV1.3-KO mice, a reduction or loss of KCNQ4, BK, and SK2 channels has been reported, accompanied by impaired efferent function in OHC [56], [57]. Our results support the idea that KCNQ4 channel function is crucial for the proper formation and maturation of efferent synapses, in line with previous reports highlighting the role of other key proteins, such as nicotinic receptors and BK channels [21], [58], [59].

Synaptophysin staining also revealed a delayed maturation process in the number and volume of MOC terminals. WT animals showed a number around 2 puncta/OHC in all tested ages, which is consistent with the finding of Barone et al., who reported an average of 2.3 boutons/OHC in C57BL/6 mice at 3, 4 and 6 postnatal weeks [60]. In *Kcnq4^-/-^*animals, this number drops significantly to below 2 puncta/OHC in 4 and 10 W animals. Similarly, aging is associated with a reduction in the number of efferent puncta, particularly in the basal and apical regions of the cochlea [41], reinforcing the idea that the structural alteration observed in our model may anticipate a progressive degenerative process. Although the number of terminals decreases with age in the KO mice, their individual volume reaches normal values at mature stages, indicating a tendency toward normalization. A similar tendency is observed for the localization of MOC terminals at this age.

The disorganization and loss of MOC terminals in the absence of KCNQ4 is consistent with reports from hearing loss mouse models associated with OHC dysfunction, such as prestin, α9 and α10 subunits of the nAChR, BK and SK2 [21], [46], [58], [59], [61]. In particular, it has been proposed that prestin provides essential signals required for the maintenance and stability of local neural networks within the OC [62].

Our results suggest that, in the absence of KCNQ4, a proportion of MOC terminals in OHC may remain, from a structural perspective, in an immature-like stage, with fewer, smaller and mislocalized efferent synapses. These terminals may later attempt to normalize, possibly to restore proper function while OHC loss is progressing from the basal to the apical turns [34]. These structural changes, together with the likely deterioration of the cochlear efferent system, correlate with the early OHC dysfunction observed through ABR and DPOAE recordings in this same model [35], suggesting that the loss of KCNQ4 leads to chronic depolarizationand compromises both the electromotile capacity of OHCs and the effectiveness of the efferent control that modulates this activity [35].

### Efferent synapses in IHC

In addition to studying efferent innervation in OHC, we also investigated the effect of KCNQ4 deletion in efferent terminals in the IHC region. As previously mentioned, the cochlea is innervated by efferent fibers that originate from two distinct groups of neurons: MOC and LOC systems. LOC fibers primarily target type I afferent dendrites that post-synapse onto IHC [63], while MOC fibers directly innervate OHCs [1]. In this study, the anti-synaptophysin signal in the IHC region was interpreted as representing LOC innervation, as these terminals are established before the onset of hearing in rodents [5]. Nevertheless, a possible contribution of MOC fibers cannot be completely ruled out [42].

Despite the changes observed in the efferent innervation of OHCs, our results indicate that the number of LOC terminals in the IHC region remains constant throughout the maturation process. However, we observed differences between genotypes at fully developed stages (4 and 10 W). The LOC terminal volume was smaller in *Kcnq4^-/-^* animals compared to their WT counterparts. Furthermore, the volume changed with age only in the *Kcnq4^+/+^* animals, reaching its maximum value at 4 W and remaining stable at 10 W. In contrast, the volume in *Kcnq4^-/-^*animals remained unaltered throughout the observed period, retaining immature characteristics. Notably, while previous studies have reported quantitative changes in the number of LOC terminals under age-related conditions [43], [64], alterations in bouton volume have not been described to our knowledge, making this observation one of the first reports of genotype-dependent structural remodeling at the level of LOC terminal size.

These results suggest that developmental or pathological changes in the LOC system may not be restricted to variations in terminal number, but could also involve subtle modifications in bouton volume, a possibility that requires further investigation.

### Elements of the efferent synapses

The observed structural disorganization of efferent terminals in KCNQ4 KO mice prompted us to investigate whether the absence of this potassium channel also dysregulates the transcriptional expression of key postsynaptic components. Our analysis revealed that the loss of KCNQ4 indeed triggers a downregulation of the key molecular machinery governing efferent inhibition, affecting both cholinergic receptors and their potassium channel effectors. The α9α10 nAChR is a heteromeric pentamer composed only by α subunits [65], that mediates inhibitory efferent cholinergic transmission in OHCs [18], [66]. As mentioned above, we observed a significant reduction in the α10 nAChR subunit expression in 4 W KO animals. Temporal changes in this subunit have been previously reported by several authors. It is known that α10 expression decreases during development, around postnatal day 13, concomitant with the loss of efferent synapses onto IHCs [14]. In contrast, in OHCs, α10 expression increases around postnatal day 10 and is maintained throughout adulthood [15], [16]. Since α10 is not expressed in mature IHCs, the decrease observed here likely reflects changes in OHCs. This reduction may result from a delay in OHC maturation, resembling a pre-postnatal day 10 stage when α10 was not yet expressed. Moreover, α10 has been shown to be required for efferent innervation of IHC [59], suggesting that its downregulation could impact efferent connectivity. Other studies have also reported an age-dependent decline in α10 expression in OHCs, but in older animals [67]. Importantly, α10 may act as a structural subunit, as it is necessary but not sufficient to sustain WT electrophysiological activity in heterologous systems and cochlear HCs [68]. Therefore, a reduction in α10 levels could compromise α9α10 nAChR function. The fact that the α9 subunit remained unchanged indicates a subunit-specific vulnerability in the regulatory mechanisms controlling nAChR expression.

Other components are the calcium-dependent potassium channels, BK and SK2. Influx of Ca^+2^ into OHCs through the nicotinic receptor leads to activation of Ca⁺^2^-dependent potassium channels, BK and SK2, which consequently results in cellular hyperpolarization through K^+^ exit via these channels. [20], [21], helping KCNQ4 to restore the membrane potential. Ca⁺^2^ influx through α9α10 nAChR is believed to trigger the release of calcium from internal cisterns located near the efferent synapse, which would activate SK2 channels [22]. BK and SK2 differs in their localization in the cochlea. BK is mainly located at the basal turn while SK2, at the apical one, and follow an opposite expression gradient from base to apex. Both are located at the basal domain of the OHC in close contact with the nAChR in the efferent synapses [21]. Similarly, to the observed for the α10 nAChR subunit, a downregulation in the expression of both potassium channels was detected in 4 W KO animals. BK channels exhibited the same number of puncta per OHC in both genotypes, but their volume was smaller and the channel localization was affected in KO animals, exhibiting a severe disruption of this precise spatial organization. We detected that the BK signal was dispersed across both the lateral and basal domains, adopting an irregular shape that worsened with age. Similar results were found for prestin by our group recently. KCNQ4 absence leads to mislocalization of prestin in OHC, extending its expression to the basal domain [35]. The BK protein analysis by Western Blot aligns with and extends these morphological observations. The significant reduction in total BK protein levels at 4 W in KO mice suggests that KCNQ4 absence may also play a role in regulating BK channel expression or stability, beyond its role in subcellular targeting. This relationship is not unexpected, as a functional interaction between both channels has been demonstrated: deletion of BK induces progressive hearing loss accompanied by a reduction in KCNQ4 expression [50]. Consequently, the absence of KCNQ4 may also alter BK expression at both the mRNA and protein levels, as supported by our data.

By 10 W, the α10 nAChR subunit, BK and SK2 mRNA levels return to wild-type baselines, despite the fact that our morphological data showed a reduction in the number of BK protein and disruption of tissue organization at this age. The age-dependent downregulation of BK protein observed in both genotypes at 10 W was unexpected and represents a novel finding. The fact that no difference in protein levels existed between WT and KO at 10 W, while a morphological disparity was evident, demonstrates that the mislocalization phenotype in the KO is not a simple consequence of reduced protein levels. Instead, it underscores the absolute necessity of the KCNQ4 function for correctly directing the available BK protein to the synapse, even when overall expression is low. A significant technical challenge arose from the inability to quantify puncta number and volume in 10 W KO animals due to the highly irregular expression pattern. This itself is a critical result, as it confirms a near-complete breakdown of discrete synaptic organization. Future ultrastructural studies using electron microscopy will be essential to determine whether physical synaptic contacts persist in the absence of molecularly defined BK plaques. Moreover, in KO animals, the number of efferent puncta decreases whereas the number of BK puncta remains unaltered, suggesting a possible uncoupling between these two proteins at the efferent synapse.

This dissociation between gene expression and protein localization, at least for BK channel, implies that developmental establishment of the efferent synapse is sensitive to the lack of KCNQ4. However, the system appears to attempt a compensatory rebound at the transcriptional level by 10 W, revealing that KCNQ4 is critical during a specific developmental window for synaptic maturation. The initial lack of key components may impair the structural integrity of the synapse. The return to normal gene expression levels at 10 W may then represent a failed attempt to repair a system that is already compromised at the subcellular localization.

In conclusion, the absence of KCNQ4 leads to a molecular response: an early significant transcriptional downregulation of key efferent postsynaptic genes (α10, BK, SK2), followed by a late transcriptional recovery. This recovery, however, is uncoupled from the protein-level (at least for BK). These results demonstrate that KCNQ4’s role extends beyond K^+^ movement function to include the regulatory orchestration of the synaptic transcriptome during a critical developmental period. In this context, the establishment of a more negative membrane potential, observed in the presence of KCNQ4, may play an important role in the transcription of efferent synaptic components, as well as in their subcellular localization.

### Calcium-binding proteins

We observed that the expression levels of two high-affinity calcium-binding proteins, calbindin and calretinin, are altered in cochlear cells. Both are known to play a critical role in shaping synaptic signaling and Ca⁺^2^ buffering, providing protection against excitotoxicity [30], [69]. Although a study in calbindin-KO animals suggested that this protein is not essential for auditory function, since ABR and DPOAE responses remained unchanged across the tested frequency range [70], it is expressed in both HCs and spiral ganglion neurons in young mice [51], but also its expression has been observed in nerve terminals and Deiters’ cells [30]. Calretinin in highly expressed in IHCs with a low expression in OHCs. In 10 W KO mice, we observed an increase in both calretinin and calbindin gene and protein expression compared to the WT counterparts and to 4 W mice. At 10 W, KO animals appear to be attempting to restore the normal localization and morphology of efferent synapses. This may explain the increased levels of these CBPs, as they could be compensating for altered intracellular calcium levels or calcium dynamics caused by the efferent defects observed in our model. Although IHC and OHC express these proteins at birth, their expression is later downregulated in OHC during the adulthood [71]. The observed transcriptional changes may originate from distinct cellular sources: the increase in calbindin transcription would be consistent with higher expression in IHCs, as suggested by the IF data, whereas the rise in calretinin transcription could be explained, at least in part, by increased expression in OHCs, also supported by the IF findings. It is interesting to note that calretinin levels are increased in OHCs from KCNQ4 animals. This CBP is more associated to IHC Ca^+2^ buffering, and this differential increase was unexpected, as more specific CBP are expressed in OHCs. However, increases in calretinin transcription levels and protein expression have been reported in OHCs during aging [72], [73].

The absence of KCNQ4 seems to delay the maturation process of the efferent innervation and the expression of these CBPs. Therefore, the increase in the expression of calbindin and calretinin may represent an adaptive change to compensate for altered calcium influx and disrupted calcium homeostasis, also suggesting that compensatory pathways may differ between IHCs and OHCs.

We also analyzed the expression of oncomodulin (Ocm), another high affinity CBP whose expression is specific to OHCs and represents the predominant CBP in the mature OHC [74]. Ocm is specifically located in basolateral domain where the efferent terminals contact the OHC [75], suggesting a direct role in regulating Ca^+2^ signals associated with synaptic activity. Ocm has been shown to contribute to the maintenance of electromotility, prevent Ca^+2^ overload, and regulate intracellular signaling during development [71], [74], [76]. In our model, Ocm expression remains unchanged at all tested ages. It is possible that the observed alterations in the efferent synapses could influence Ocm expression, but only at the protein level. Further studies will be required to explore this possibility.

Based on this evidence, a hypothesis could be proposed regarding the differential function of CBPs in each sensory cell type. The increase in calbindin in IHCs could help buffer changes in intracellular Ca⁺^2^ levels associated with their depolarization, and, considering the low basal expression of KCNQ4 in these cells, this enhanced Ca⁺^2^-buffering capacity may prevent depolarization from reaching cytotoxic levels, thereby preserving cell integrity and providing greater resistance to channel dysfunction. In contrast, although calretinin levels rise in OHCs, this increase is more limited and occurs in a context in which the major CBP (Ocm) remains unchanged. This suggests that the elevation in calretinin represents a compensatory attempt to counteract the consequences in calcium homeostasis that the sustained depolarization that characterizes OHCs in the absence of KCNQ4 generates. Alternatively, the changes in CBP expression may be related to their distinct subcellular localization. Ocm is mainly associated with regions close to the plasma membrane, whereas calretinin in largely cytosolic. Thus, these expression changes may reflect differences in the subcellular domains in which intracellular calcium dynamics are affected. Overall, this hypothesis therefore proposes that the differential vulnerability between OHCs and IHCs may arise from a combination of the magnitude of chronic depolarization and the relative ability of each cell type to upregulate its Ca⁺^2^ -buffering mechanisms.

## Conclusion

Taken together, our current and previous findings [34], [35], in combination with prior work demonstrating chronic depolarization in the absence of KCNQ4 [27], suggest that many of the alterations observed in *Kcnq4^−/−^*animals arise from a sustained disruption of the membrane potential in HCs. This electrical imbalance perturbs ionic gradients and signaling dynamics, triggering a degenerative cascade that compromises tissue organization. Such effects align with the framework of bioelectricity, a regulatory level in which ionic gradients, membrane potentials, and intercellular electrical networks act as instructive circuits governing tissue architecture, identity, and morphogenesis [77]–[79]. From this perspective, proper membrane voltage is essential for differentiation, adhesion, migration, and maturation, particularly in highly specialized tissues such as the OC [78].

Our results further indicate that KCNQ4 is required for the correct subcellular positioning and long-term maintenance of presynaptic SYN terminals and postsynaptic BK channels at OHC efferent synapses. The observed mislocalization, reduction in boutons volume, and progressive synaptic disorganization are compatible with a role for KCNQ4 in mechanisms related to cytoskeletal anchoring, trafficking, or the stabilization of synaptic organizational complexes. Importantly, the dissociation between transcript levels and synaptic protein localization points to post-transcriptional regulatory processes that may be particularly sensitive to changes in the cellular electrical environment.

Overall, our findings support a model in which chronic alterations in membrane potential, as described in the absence of KCNQ4, may interfere with the bioelectric conditions required for proper maturation and maintenance of cochlear efferent synapses. While this hypothesis remains to be directly tested, it raises the possibility that bioelectric regulation represents a previously underappreciated layer of cochlear homeostasis. Future studies addressing membrane voltage dynamics and their causal relationship with synaptic development will be essential to validate this model and to evaluate whether restoring electrical balance could hold therapeutic relevance for DFNA2-associated hearing loss.

## Statements and Declarations

## Acknowledgements

Our special thanks to Lic. Karen Schweitzer, for helping with WB experiments, and Prof. Thomas Jentsch (Max-Delbrück- Centrum für Molekulare Medizin and Leibniz-Institut für Molekulare Pharmakologie, Berlin, Germany) for generously providing *Kcnq4^−/−^* animals.

## Funding

This work was supported by grants from Agencia Nacional de Promoción Científica y Tecnológica (ANPCyT) (PICT 2021-0145), CONICET (PIP 0091) and UNS (PGI 24/B262) to GS; and also, by grants from ANPCyT (PICT 2021-0545), CONICET (PIBAA 0769) and UNS (PGI 24/ZB98) to LD. IO had an advanced student fellowship from the UNS.

### Author Contributions

Ezequiel Rías: Conceptualization, Formal Analysis, Investigation, Methodology, Software, Resources, Validation, Visualization, Writing- original draft, Writing- review and editing.

Ingrid Ouwerkerk: Formal Analysis, Investigation, Methodology, Software, Resources, Validation.

Guillermo Spitzmaul: Conceptualization, Funding Acquisition, Investigation, Project Administration, Resources, Supervision, Visualization, Writing- original draft, Writing-review and editing.

Leonardo Dionisio: Conceptualization, Formal Analysis, Funding Acquisition, Investigation, Methodology, Project Administration, Resources, Supervision, Validation, Visualization, Writing- original draft, Writing- review and editing.

### Data Availability

The datasets generated during and/or analysed during the current study are not publicly available due to a large amount of data but are available rom the corresponding author on reasonable request.

### Ethics Approval

The experimental protocol followed in this study was approved by the Council for Care and Use of Experimental Animals (CICUAE, protocol no. 083/2016) of the Universidad Nacional del Sur (UNS), whose requirements are strictly based on the European Parliament and Council of the European Union directives (2010/63/EU).

## References

[1] J. J. Guinan, “Olivocochlear efferents: Their action, effects, measurement and uses, and the impact of the new conception of cochlear mechanical responses,” Hear. Res., vol. 362, pp. 38–47, 2018, doi: 10.1016/j.heares.2017.12.012.

[2] P. A. Fuchs, “Synaptic transmission at vertebrate hair cells,” Curr. Opin. Neurobiol., vol. 6, no. 4, 1996, doi: 10.1016/S0959-4388(96)80058-4.

[3] J. Guinan J. J., “Olivocochlear efferents: anatomy, physiology, function, and the measurement of efferent effects in humans,” Ear Hear., vol. 27, no. 6, pp. 589–607, 2006.

[4] A. A. Sitko and and L. V. G. Michelle M. Franka, Gabriel E. Romeroa, Mackenzie Hunta, “Lateral olivocochlear neurons modulate cochlear responses to noise exposure,” PNAS, vol. 122, pp. 1–12, 2025, doi: 10.1073/pnas.

[5] A. V. Bulankina and T. Moser, “Neural circuit development in the mammalian cochlea,” Physiology, vol. 27, no. 2, pp. 100–112, 2012, doi: 10.1152/physiol.00036.2011.

[6] J. K. Morris et al., “A disorganized innervation of the inner ear persists in the absence of ErbB2,” Brain Res., vol. 1091, no. 1, pp. 186–199, 2006, doi: 10.1016/j.brainres.2006.02.090.

[7] A. Shnerson, C. Devigne, and R. Pujol, “Age-related changes in the C57BL/6J mouse cochlea. II. Ultrastructural findings,” Dev. Brain Res., vol. 2, no. 1, pp. 77–88, 1981, doi: 10.1016/0165-3806(81)90060-2.

[8] M. R. Emmerling, H. M. Sobkowicz, C. V. Levenick, G. L. Scott, S. M. Slapnick, and J. E. Rose, “Biochemical and morphological differentiation of acetylcholinesterasepositive efferent fibers in the mouse cochlea,” J. Electron Microsc. Tech., vol. 15, no. 2, pp. 123–143, 1990, doi: 10.1002/jemt.1060150205.

[9] M. Knipper, U. Zimmermann, K. Rohbock, I. Köpschall, and H. P. Zenner, “Synaptophysin and Gap-43 proteins in efferent fibers of the inner ear during postnatal development,” Dev. Brain Res., vol. 89, no. 1, pp. 73–86, 1995, doi: 10.1016/0165-3806(95)00113-R.

[10] A. B. Elgoyhen, D. S. Johnson, J. Boulter, D. E. Vetter, and S. Heinemann, “α9: An acetylcholine receptor with novel pharmacological properties expressed in rat cochlear hair cells,” Cell, vol. 79, no. 4, pp. 705–715, 1994, doi: 10.1016/0092-8674(94)90555-X.

[11] I. Roux, E. Wersinger, J. M. McIntosh, P. A. Fuchs, and E. Glowatzki, “Onset of cholinergic efferent synaptic function in sensory hair cells of the rat cochlea,” J. Neurosci., vol. 31, no. 42, pp. 15092–15101, 2011, doi: 10.1523/JNEUROSCI.2743-11.2011.

[12] K. S. Cole and D. Robertson, “Early efferent innervation of the developing rat cochlea studied with a carbocyanine dye,” Brain Res., vol. 575, no. 2, pp. 223–230, 1992, doi: 10.1016/0006-8993(92)90083-L.

[13] R. Pujol, M. Lavigne-Rebillard, and M. Lenoir, “Development of Sensory and Neural Structures in the Mammalian Cochlea,” in Development of the Auditory System, 9th ed., R. R. Rubel, E.W., Popper, A.N., Fay, Ed. New York: Springer, New York, NY, 1998, pp. 146–192. doi: 10.1007/978-1-4612-2186-9_4.

[14] E. Katz et al., “Developmental regulation of nicotinic synapses on cochlear inner hair cells,” J. Neurosci., vol. 24, no. 36, pp. 7814–7820, 2004, doi: 10.1523/JNEUROSCI.2102-04.2004.

[15] A. B. Elgoyhen, D. E. Vetter, E. Katz, C. V. Rothlin, S. F. Heinemann, and J. Boulter, “α10: A determinant of nicotinic cholinergic receptor function in mammalian vestibular and cochlear mechanosensory hair cells,” Proc. Natl. Acad. Sci. U. S. A., vol. 98, no. 6, pp. 3501–3506, 2001, doi: 10.1073/pnas.051622798.

[16] B. J. Morley and D. D. Simmons, “Developmental mRNA expression of the α10 nicotinic acetylcholine receptor subunit in the rat cochlea,” Dev. Brain Res., vol. 139, no. 1, pp. 87–96, 2002, doi: 10.1016/S0165-3806(02)00514-X.

[17] P. A. Fuchs and A. M. Lauer, “Efferent Inhibition of the Cochlea,” Cold Spring Harb. Perspect. Med., vol. 9, no. 5, pp. 1–15, 2019, doi: 10.1101/cshperspect.a033530.

[18] M. E. Gómez-Casati, P. A. Fuchs, A. B. Elgoyhen, and E. Katz, “Biophysical and pharmacological characterization of nicotinic cholinergic receptors in rat cochlear inner hair cells,” J. Physiol., vol. 566, no. 1, pp. 103–118, 2005, doi: 10.1113/jphysiol.2005.087155.

[19] N. Weisstaub, D. E. Vetter, A. Belén Elgoyhen, and E. Katz, “The α9α10 nicotinic acetylcholine receptor is permeable to and is modulated by divalent cations,” Hear. Res., vol. 167, no. 1–2, pp. 122–135, 2002, doi: 10.1016/S0378-5955(02)00380-5.

[20] K. N. Rohmann, E. Wersinger, J. P. Braude, S. J. Pyott, and P. A. Fuchs, “Activation of BK and SK channels by efferent synapses on outer hair cells in high-frequency regions of the rodent cochlea,” J. Neurosci., vol. 35, no. 5, pp. 1821–1830, 2015, doi: 10.1523/JNEUROSCI.2790-14.2015.

[21] S. F. Maison, S. J. Pyott, A. L. Meredith, and M. C. Liberman, “Olivocochlear suppression of outer hair cells in vivo: Evidence for combined action of BK and SK2 channels throughout the cochlea,” J. Neurophysiol., vol. 109, no. 6, pp. 1525–1534, 2013, doi: 10.1152/jn.00924.2012.

[22] P. A. Fuchs, M. Lehar, and H. Hiel, “Ultrastructure of Cisternal Synapses on Outer Hair Cells of the Mouse Cochlea,” J Comp Neurol., pp. 717–729, 2014, doi: 10.1146/annurev-biophys-062420-081842.The.

[23] B. Fakler and J. P. Adelman, “Control of KCa Channels by Calcium Nano/Microdomains,” Neuron, vol. 59, no. 6, pp. 873–881, 2008, doi: 10.1016/j.neuron.2008.09.001.

[24] J. D. Goutman, A. B. Elgoyhen, and M. E. Gomez-Casati, “Cochlear hair cells: the sound-sensing machines,” Physiol. Behav., 2015, doi: 10.1016/j.febslet.2015.08.030.Cochlear.

[25] S. J. Pyott and R. K. Duncan, BK Channels in the Vertebrate Inner Ear, 1st ed., vol. 128. Elsevier Inc., 2016. doi: 10.1016/bs.irn.2016.03.016.

[26] E. Wersinger, W. J. McLean, P. A. Fuchs, and S. J. Pyott, “BK channels mediate cholinergic inhibition of high frequency cochlear hair cells,” PLoS One, vol. 5, no. 11, 2010, doi: 10.1371/journal.pone.0013836.

[27] T. Kharkovets et al., “Mice with altered KCNQ4 K+ channels implicate sensory outer hair cells in human progressive deafness,” EMBO J., vol. 25, no. 3, 2006, doi: 10.1038/sj.emboj.7600951.

[28] J. H. Rim, J. Y. Choi, J. Jung, and H. Y. Gee, “Activation of kcnq4 as a therapeutic strategy to treat hearing loss,” Int. J. Mol. Sci., vol. 22, no. 5, pp. 1–12, 2021, doi: 10.3390/ijms22052510.

[29] E. Katz and A. B. Elgoyhen, “Short-term plasticity and modulation of synaptic transmission at mammalian inhibitory cholinergic olivocochlear synapses,” Front. Syst. Neurosci., vol. 8, no. DEC, pp. 1–14, 2014, doi: 10.3389/fnsys.2014.00224.

[30] C. M. Hackney, S. Mahendrasingam, A. Penn, and R. Fettiplace, “The concentrations of calcium buffering proteins in mammalian cochlear hair cells,” J. Neurosci., vol. 25, no. 34, pp. 7867–7875, 2005, doi: 10.1523/JNEUROSCI.1196-05.2005.

[31] P. Delmas and D. A. Brown, “PATHWAYS MODULATING NEURAL KCNQ / M C Kv7 C POTASSIUM CHANNELS,” vol. 6, no. November, pp. 850–862, 2005, doi: 10.1038/nrn1785.

[32] J. Taranda et al., “A point mutation in the hair cell nicotinic cholinergic receptor prolongs cochlear inhibition and enhances noise protection,” PLoS Biol., vol. 7, no. 1, 2009, doi: 10.1371/journal.pbio.1000018.

[33] L. E. Boero et al., “Preventing presbycusis in mice with enhanced medial olivocochlear feedback,” Proc. Natl. Acad. Sci. U. S. A., vol. 117, no. 21, 2020, doi: 10.1073/pnas.2000760117.

[34] C. Carignano, E. P. Barila, E. I. Rías, L. Dionisio, E. Aztiria, and G. Spitzmaul, “Inner Hair Cell and Neuron Degeneration Contribute to Hearing Loss in a DFNA2-Like Mouse Model,” Neuroscience, vol. 410, 2019, doi: 10.1016/j.neuroscience.2019.05.012.

[35] E. Rías et al., “Insights into early cochlear damage induced by potassium channel deficiency,” Biochim. Biophys. Acta - Mol. Cell Res., vol. 1872, no. 8, p. 120030, 2025, doi: 10.1016/j.bbamcr.2025.120030.

[36] Guillermo Spitzmaul, E. Rías, and L. Dionisio, “Potential Mechanisms of Hearing Loss Due to Impaired Potassium Circulation in the Organ of Corti,” in *Hearing Loss - Diagnosis*, Management and Future Challenges, 2023.

[37] A. Myint et al., “Large-scale Phenotyping of Noise-Induced Hearing Loss in 100 Strains of Mice,” pp. 113–120, 2016, doi: 10.1016/j.heares.2015.12.006.Large-scale.

[38] D. R. Trune, J. B. Kempton, and C. Mitchell, “Auditory function in the C3H/HeJ and C3H/HeSnJ mouse strains,” Hear. Res., vol. 96, no. 1–2, pp. 41–45, 1996, doi: 10.1016/0378-5955(96)00017-2.

[39] J. Y. Jeng et al., “Age-related changes in the biophysical and morphological characteristics of mouse cochlear outer hair cells,” J. Physiol., vol. 598, no. 18, pp. 3891–3910, 2020, doi: 10.1113/JP279795.

[40] S. F. Maison, A. E. Luebke, M. C. Liberman, and J. Zuo, “Efferent protection from acoustic injury is mediated via α9 nicotinic acetylcholine receptors on outer hair cells,” J. Neurosci., vol. 22, no. 24, pp. 10838–10846, 2002, doi: 10.1523/jneurosci.22-24-10838.2002.

[41] N. M. Dörje et al., “Age-related alterations in efferent medial olivocochlear-outer hair cell and primary auditory ribbon synapses in CBA/J mice,” Front. Cell. Neurosci., vol. 18, no. June, pp. 1–16, 2024, doi: 10.3389/fncel.2024.1412450.

[42] Y. Hua et al., “Electron Microscopic Reconstruction of Neural Circuitry in the Cochlea,” Cell Rep., vol. 34, no. 1, p. 108551, 2021, doi: 10.1016/j.celrep.2020.108551.

[43] F. Steenken, A. Pektaş, and C. Köppl, “Age-related changes in olivocochlear efferent innervation in gerbils,” Front. Synaptic Neurosci. , vol. 16, no. June, pp. 1–17, 2024, doi: 10.3389/fnsyn.2024.1422330.

[44] K. J. Livak and T. D. Schmittgen, “Analysis of relative gene expression data using real-time quantitative PCR and the 2-ΔΔCT method,” Methods, vol. 25, no. 4, pp. 402–408, 2001, doi: 10.1006/meth.2001.1262.

[45] T. Bayasgalan et al., “Alteration of Mesopontine Cholinergic Function by the Lack of KCNQ4 Subunit,” Front. Cell. Neurosci., vol. 15, no. July, pp. 1–16, 2021, doi: 10.3389/fncel.2021.707789.

[46] S. Takahashi et al., “Prestin contributes to membrane compartmentalization and is required for normal innervation of outer hair cells,” Front. Cell. Neurosci., vol. 12, no. July, pp. 1–11, 2018, doi: 10.3389/fncel.2018.00211.

[47] M. V. Bartolome, P. Zuluaga, F. Carricondo, and P. Gil-Loyzaga, “Immunocytochemical detection of synaptophysin in C57BL/6 mice cochlea during aging process,” Brain Res. Rev., vol. 60, no. 2, pp. 341–348, 2009, doi: 10.1016/j.brainresrev.2009.02.001.

[48] L. G. Vattino, C. Wedemeyer, A. B. Elgoyhen, and E. Katz, “Functional Postnatal Maturation of the Medial Olivocochlear Efferent–Outer Hair Cell Synapse,” J. Neurosci., vol. 40, no. 25, pp. 4842–4857, 2020, doi: 10.1523/JNEUROSCI.2409-19.2020.

[49] R. Pujol and M. Lavigne-Rebillard, “Development and plasticity of the human auditory system,” *A Textb*. Audiol. Med., no. February, pp. 147–156, 2002, doi: 10.1201/b14730-12.

[50] L. Rüttiger et al., “Deletion of the Ca2+-activated potassium (BK) α-subunit but not the BKβ1-subunit leads to progressive hearing loss,” Proc. Natl. Acad. Sci. U. S. A., vol. 101, no. 35, pp. 12922–12927, 2004, doi: 10.1073/pnas.0402660101.

[51] W. Liu, H. Chen, X. Zhu, and H. Yu, “Expression of Calbindin-D28K in the Developing and Adult Mouse Cochlea,” vol. 70, no. 8, 2022, doi: 10.1369/00221554221119543.

[52] X. Zhang, H. Wang, and Q. Wang, “Progression of KCNQ4 related genetic hearing loss: a narrative review,” J. Bio-X Res., vol. 4, no. 4, pp. 151–157, 2021, doi: 10.1097/JBR.0000000000000112.

[53] L. M. Dominguez and K. M. Dodson, “Genetics of hearing loss,” Oxford Handb. Audit. Sci. Ear, pp. 97–104, 2012, doi: 10.1093/oxfordhb/9780199233397.013.0014.

[54] H. Winter et al., “Thyroid hormone receptors TRα1 and TRβ differentially regulate gene expression of Kcnq4 and prestin during final differentiation of outer hair cells,” J. Cell Sci., vol. 119, no. 14, pp. 2975–2984, 2006, doi: 10.1242/jcs.03013.

[55] M. C. Perez-Flores et al., “Cooperativity of Kv7.4 channels confers ultrafast electromechanical sensitivity and emergent properties in cochlear outer hair cells,” Sci. Adv., vol. 6, no. 15, pp. 1–11, 2020, doi: 10.1126/sciadv.aba1104.

[56] J. Y. Jeng et al., “Hair cell maturation is differentially regulated along the tonotopic axis of the mammalian cochlea,” J. Physiol., vol. 598, no. 1, pp. 151–170, 2020, doi: 10.1113/JP279012.

[57] J. Ashmore, “A maturing view of cochlear calcium,” J. Physiol., vol. 598, no. 1, pp. 7–8, 2020, doi: 10.1113/JP279215.

[58] and D. E. V. Vidya Murthy, Stéphane F. Maison, Julián Taranda, Nadeem Haque, Chris T. Bond, A. Belén Elgoyhen, John P. Adelman, M. Charles Liberman, “SK2 channels are required for function and long-term survival of efferent synapses on mammalian outer hair cells,” Mol. Cell Neurosci., vol. 40, no. 1, pp. 39–49, 2009, doi: 10.1016/j.mcn.2008.08.011.SK2.

[59] D. E. Vetter et al., “The α10 nicotinic acetylcholine receptor subunit is required for normal synaptic function and integrity of the olivocochlear system,” Proc. Natl. Acad. Sci. U. S. A., vol. 104, no. 51, pp. 20594–20599, 2007, doi: 10.1073/pnas.0708545105.

[60] C. M. Barone et al., “Altered cochlear innervation in developing and mature naked and Damaraland mole rats,” J. Comp. Neurol., vol. 527, no. 14, pp. 2302–2316, 2019, doi: 10.1002/cne.24682.

[61] D. E. Vetter et al., “Role of α9 nicotinic ACh receptor subunits in the development and function of cochlear efferent innervation,” Neuron, vol. 23, no. 1, pp. 93–103, 1999, doi: 10.1016/S0896-6273(00)80756-4.

[62] J. Zheng, W. Shen, D. Z. Z. He, K. B. Long, L. D. Madison, and P. Dallos, “Prestin is the motor protein of cochlear outer hair cells,” Nature, vol. 405, no. 6783, pp. 149–155, 2000, doi: 10.1038/35012009.

[63] J. Ruel et al., “Physiology, pharmacology and plasticity at the inner hair cell synaptic complex,” Hear. Res., vol. 227, no. 1–2, pp. 19–27, 2007, doi: 10.1016/j.heares.2006.08.017.

[64] A. M. Lauer, P. A. Fuchs, D. K. Ryugo, and Howard W. Francis, “Efferent synapses return to inner hair cells in the aging cochlea,” Neurobiol Aging, vol. 33, no. 12, pp. 2892–2902, 2012, doi: 10.1016/j.neurobiolaging.2012.02.007.Efferent.

[65] A. B. Elgoyhen, “The α9α10 nicotinic acetylcholine receptor: a compelling drug target for hearing loss?,” Expert Opin. Ther. Targets, vol. 26, no. 3, pp. 291–302, 2022, doi: 10.1080/14728222.2022.2047931.

[66] A. B. Elgoyhen and E. Katz, “The Efferent Medial Olivocochlear-Hair Cell Synapse,” J Physiol Paris, vol. 106, no. 1–2, pp. 47–56, 2012, doi: 10.1016/j.jphysparis.2011.06.001.The.

[67] H. Liu et al., “Molecular and cytological profiling of biological aging of mouse cochlear inner and outer hair cells,” Cell Rep., vol. 39, no. 2, 2022, doi: 10.1016/j.celrep.2022.110665.

[68] B. J. Morley, D. F. Dolan, K. K. Ohlemiller, and D. D. Simmons, “Generation and characterization of α9 and α10 nicotinic acetylcholine receptor subunit knockout mice on a C57BL/6J background,” Front. Neurosci., vol. 11, no. SEP, pp. 1–15, 2017, doi: 10.3389/fnins.2017.00516.

[69] Y.-W. S. and E. Y. Kushal Sharma, “Differential Expression of Ca 2 + -buffering Protein Calretinin in Cochlear Afferent FibersC: A Possible Link to Vulnerability to Traumatic Noise,” vol. 27, no. 5, pp. 397–407, 2018.

[70] L. Airaksinen, J. Virkkala, A. Aarnisalo, M. Meyer, J. Ylikoski, and M. S. Airaksinen, “Lack of calbindin-D28k does not affect hearing level or survival of hair cells in acoustic trauma,” Orl, vol. 62, no. 1, pp. 9–12, 1999, doi: 10.1159/000027708.

[71] B. Tong, A. J. Hornak, S. F. Maison, K. K. Ohlemiller, M. C. Liberman, and D. D. Simmons, “Oncomodulin, an EF-hand Ca2+ buffer, is critical for maintaining cochlear function in mice,” J. Neurosci., vol. 36, no. 5, pp. 1631–1635, 2016, doi: 10.1523/JNEUROSCI.3311-15.2016.

[72] M. Zhang, W. Liu, D. Ding, and R. Salvi, “Pifithrin-α supresses p53 and protects cochlear and vestibular hair cells from cisplatin-induced apoptosis,” Neuroscience, vol. 120, no. 1, pp. 191–205, 2003, doi: 10.1016/S0306-4522(03)00286-0.

[73] A. O. Sodero et al., “Phytosterols reverse antiretroviral-induced hearing loss, with potential implications for cochlear aging,” PLoS Biol., vol. 21, no. 8 August, pp. 1–22, 2023, doi: 10.1371/journal.pbio.3002257.

[74] K. E. Murtha et al., “Oncomodulin (OCM) uniquely regulates calcium signaling in neonatal cochlear outer hair cells,” Cell Calcium, vol. 105, no. June, p. 102613, 2022, doi: 10.1016/j.ceca.2022.102613.

[75] and A. J. H. Dwayne D. Simmons, Benton Tong, Angela D. Schrader, “Oncomodulin Identifies Different Hair Cell Types in the Mammalian Inner Ear,” Natl. Inst. Heal., vol. 518, no. 18, pp. 3785–3802, 2010, doi: 10.1002/cne.22424.Oncomodulin.

[76] Y. Yang et al., “Oncomodulin regulates spontaneous calcium signalling and maturation of afferent innervation in cochlear outer hair cells,” J. Physiol., vol. 601, no. 19, pp. 4291–4308, 2023, doi: 10.1113/JP284690.

[77] M. Levin, “Bioelectric signaling: Reprogrammable circuits underlying embryogenesis, regeneration, and cancer,” Cell, vol. 184, no. 8, pp. 1971–1989, 2021, doi: 10.1016/j.cell.2021.02.034.

[78] L. F. George and E. A. Bates, “Mechanisms Underlying Influence of Bioelectricity in Development,” Front. Cell Dev. Biol., vol. 10, no. February, pp. 1–17, 2022, doi: 10.3389/fcell.2022.772230.

[79] C. O. Nunes and E. H. Barriga, “Bioelectricity in Morphogenesis,” Annu. Rev. Cell Dev. Biol., vol. 41, no. 1, pp. 187–208, 2025, doi: 10.1146/annurev-cellbio-101323-032747.

